# Interacting Learning Processes during Skill Acquisition: Learning to control with gradually changing system dynamics

**DOI:** 10.1101/135095

**Authors:** Nicolas Ludolph, Martin A. Giese, Winfried Ilg

**Affiliations:** Section Computational Sensomotorics, Department of Cognitive Neurology, Hertie Institute for Clinical Brain Research, and Centre for Integrative Neuroscience, University of Tübingen, Germany; International Max-Planck Research School for Cognitive and Systems Neuroscience, University of Tübingen, Germany

**Keywords:** motor learning, skill acquisition, motor adaptation, internal forward models, predictive control, action timing

## Abstract

There is increasing evidence that sensorimotor learning under real-life conditions relies on a composition of several learning processes. Nevertheless, most studies examine learning behaviour in relation to one specific learning mechanism. In this study, we examined the interaction between reward-based skill acquisition and motor adaptation to changes of object dynamics. Thirty healthy subjects, split into two groups, acquired the skill of balancing a pole on a cart in virtual reality. In one group, we gradually increased the gravity, making the task easier in the beginning and more difficult towards the end. In the second group, subjects had to acquire the skill on the maximum, most difficult gravity level. We hypothesized that the gradual increase in gravity during skill acquisition supports learning despite the necessary adjustments to changes in cart-pole dynamics. We found that the gradual group benefits from the slow increment, although overall improvement was interrupted by the changes in gravity and resulting system dynamics, which caused short-term degradations in performance and timing of actions. In conclusion, our results deliver evidence for an interaction of reward-based skill acquisition and motor adaptation processes, which indicates the importance of both processes for the development of optimized skill acquisition schedules.

## INTRODUCTION

Sensorimotor learning is a term commonly used to capture the variety of neural changes taking place (i) during the acquisition of a new motor skill or (ii) during the adaptation of movements to changed circumstances such as changes of the environment. Most extensively, motor adaptation to external perturbations has been studied over the last decades in experimental paradigms like visuomotor rotation^1,2^ and velocity-dependent force-fields^3,4^. The underlying mechanism is considered to be a recalibration of internal forward models, which predict the sensory consequences of the motor action^5^,^6^. During the adaptation process, these internal forward models are re-calibrated based on a sensory prediction error^7^ which is the difference between the internal forward model prediction and the actual sensory outcome.

In recent years, a particular interest has emerged in studies which compare the sudden application of perturbations to gradually induced perturbations^8,9^, as well as the influence of reward-based feedback on the adaptation process^10-12^. In contrast to the described motor adaptation paradigms, motor skill acquisition describes the expansion of the motor repertoire when faced with completely new demands, such as learning for the first time how to ride a bicycle, monocycle or acquiring a new sports skill^13-15^.

Although studies in sports science have been examining skill acquisition for many years on a descriptive level^14,16,17^, computational studies investigating the underlying control mechanisms have mainly been restricted to simplified experimental paradigms. Examples for such paradigms are the learning of finger-tapping sequences^18^, visually-guided hand movement trajectories^15^, simplified virtual-reality versions of moving a cup^19^, bouncing a ball^20,21^ or playing skittles^22^. The focus of these studies is mainly the quantitative analysis of the execution performance and performance variability within the skill acquisition process13. However, processes of acquiring or adjusting internal models and specific control mechanisms like predictive control have not been the addressed in this context.

Predictive control based on forward models is suggested to play an important role in many dynamic skills like bouncing or catching a ball^17^ and performing fast goal directed movements^23^–^25^. More generally speaking, skilled motor behaviour is suggested to rely on accurate predictive models of both our own body and tools we interact with26. Indeed, studies have collected increasing evidence that the brain acquires and uses an internal model that encodes the physical properties of our limbs^27^, environment and manipulated objects^28–31^. Thus, internal forward models of new tools or objects have to be acquired during skill acquisition and have to be adjusted to changes of body dynamics during development or to external changes when the dynamics of the object change^32^.

The dynamics of external objects change for instance frequently during skill acquisition when different objects or sport devices are used as part of the training, such as different tennis rackets^33–35^. In the beginning of the learning process devices are used which are easier to handle and thus lead earlier to successful behaviour. These moments of success and resultant motivation are known to be important factors in motor learning and have been reported in several studies on motor adaptation or learning in sports^36–38^. On the other hand, at the transition to another device, the subject has to adapt the control behaviour to the dynamics of the new device within the process of skill acquisition. Thus, two learning processes, one driven by reward (skill acquisition) and the other by sensory prediction error (motor adaptation), shape the behaviour concurrently. In this study, we want to investigate the influence of interleaved adaptation phases during skill acquisition by examining subjects’ behaviour in the virtual cart-pole balancing task while manipulating the gravity. Increments in gravity do not only change the dynamic behaviour of the system but also the difficulty to control it, allowing us to study the interaction of reward-based and error-based learning. Cart-pole balancing has been studied in the context of reinforcement learning as a benchmark for computational algorithms^39^, as model for human balance control^40–44^ and in the context of internal forward models^30,44^. We investigate the implications of gradually increasing the task complexity (i) on the skill acquisition and (ii) in terms of the need to adapt to repetitively changed cart-pole dynamics. The hypothesis under investigation is that a gradual increase in task complexity leads to improved learning due to earlier and more frequent positive reward, which is however disturbed by short interleaved adaptation phases caused by the gradually changing cart-pole dynamics.

## RESULTS

### Experimental Design

Participants had to learn the control of a physically simulated cart-pole system (Fig 1A, see Methods for details). Gravity forces the pole to rotate downwards when not being perfectly upright. Thus, in the simulation, the complexity of the control task can gradually manipulated by the adjustment of the simulated gravity. Higher gravity leads to a faster falling pole and to a more complex control problem. Participants controlled the car by applying virtual force using a haptic input device (Fig 1B). The goal was to keep the pole upright. Specifically, the pole has to remain within the green circular segment (±60 degree, Fig 1A) while the cart must not leave the track (±5m). A trial is considered as successful, if balance was maintained for 30 seconds without violating the two given constrains.

We examined two groups of subjects corresponding to different experimental conditions (Fig 1C): (i) gradual gravity (GG), starting with a low gravitational constant of g_0_=1.0m/s^2^, the virtual gravity has been increased in relation to the current performance up to the maximum of g_max_=3.5m/s^2^. Specifically, the gravitational constant was increased by 0.1m/s^2^ after every successful trial (performance-dependent increase in gravity). (ii) Constant gravity (CG), in this group the gravity has been kept constant on the maximum of g_max_=3.5m/s^2^ from the beginning. Notice, that the gravitational constant was never decreased and that every subject was exposed to an individual gravity profile over the course of the experiment, due to the performance-dependent increase. Subjects in both groups interacted for 90 minutes with the cart-pole environment, while the number of trials during this time was not limited.

**Fig 1.**
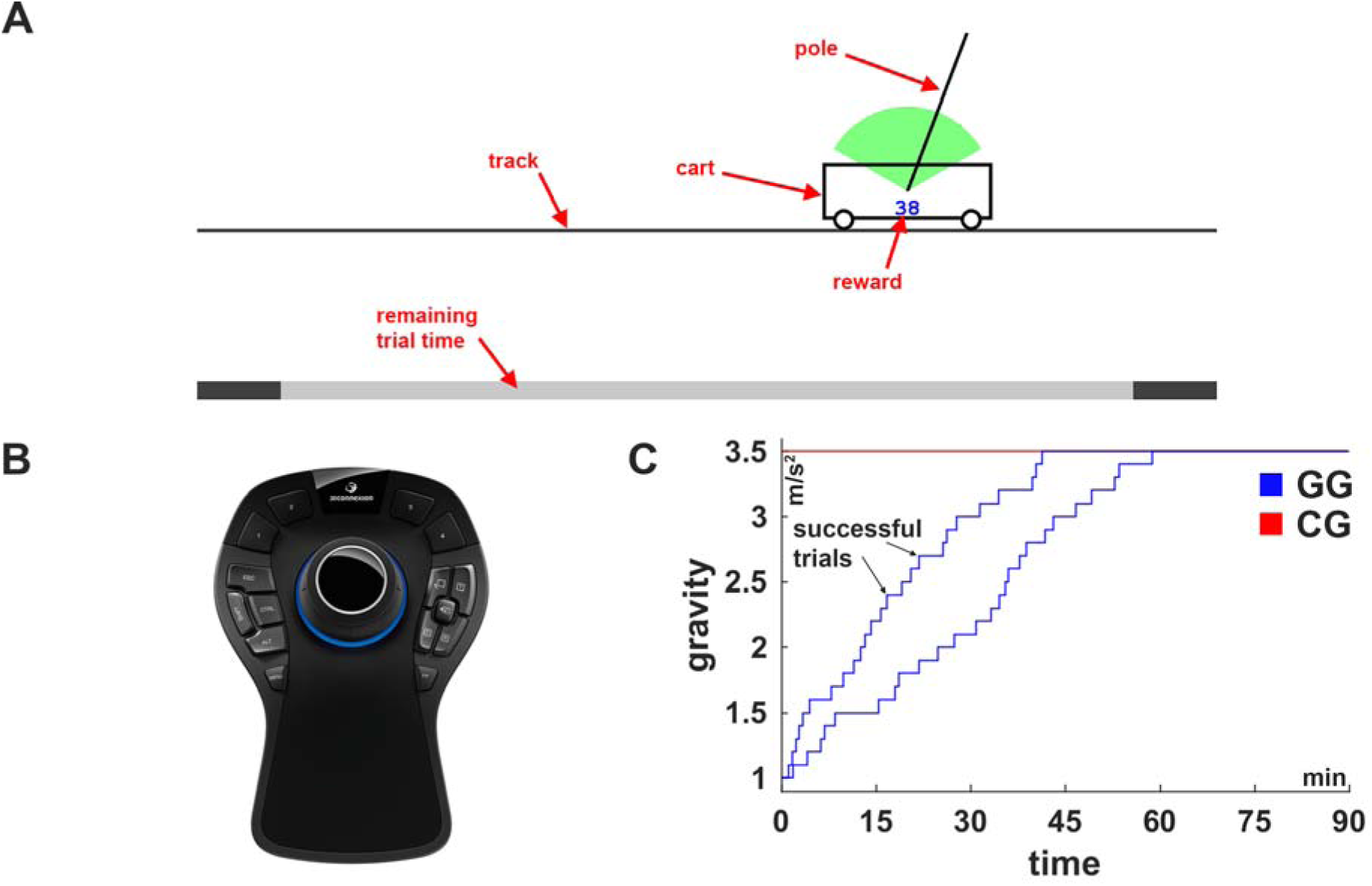
Illustrations of the cart-pole system, the used input device and experimental conditions. Cart-pole system. (B) Input device. The knob of the input device can be shifted and rotated into all directions. The left-right translation was used to control the virtual force, which is applied to the cart from either side. (C) Experimental conditions: gradual gravity (GG) and constant gravity (CG). For the condition GG, the course of the gravity is shown for two representative subjects to illustrate the individual, performance-dependent increase. Gravity was increased after every successful balancing attempt.

## Task-Performance

Task performance, measured by the trial length, is shown in Fig 2 as running average over the course of the experiment for both groups. Eventhough the performance looks almost constant for the gradual group (Fig 2A), keep in mind that the gravitational constant was increased after every successful trial making the task more difficult over time. In order to account for this influence, we also examined the improvement based on the normalized trial length T/T_0_ (Fig 2B), indicating that both groups improve monotonically.

Analysis of gender (Wilcoxon rank sum, **CG: p=0.28, GG: p=1.0**) or age as influential factors on the improvement in task performance (Pearson’s linear correlation coefficient, **CG: rho=-0.25, p=0.36, GG: rho=-0.28, p=0.31**)did not yield any significant correlations within the two groups. Comparing the average task performance within the first 5 minutes of the experiment, reveals significantly higher performance for subjects in condition GG (Wilcoxon rank sum, **p<0.001**). As the only difference between the groups is the difference in gravity during this phase, this result verifies our expectation about strong influence of the gravity on the task difficulty. The difference in task performance between the groups however persists throughout the experiment. Finally, subjects in the gradual group are also significantly better at the end (last 5 minutes) of the experiment (trial length, Wilcoxon rank sum, **p<0.05**), despite the fact that the gravity is identical for both groups at this point. The average course of the gravitational constant is shown in Fig 2 D.

**Fig 2.**
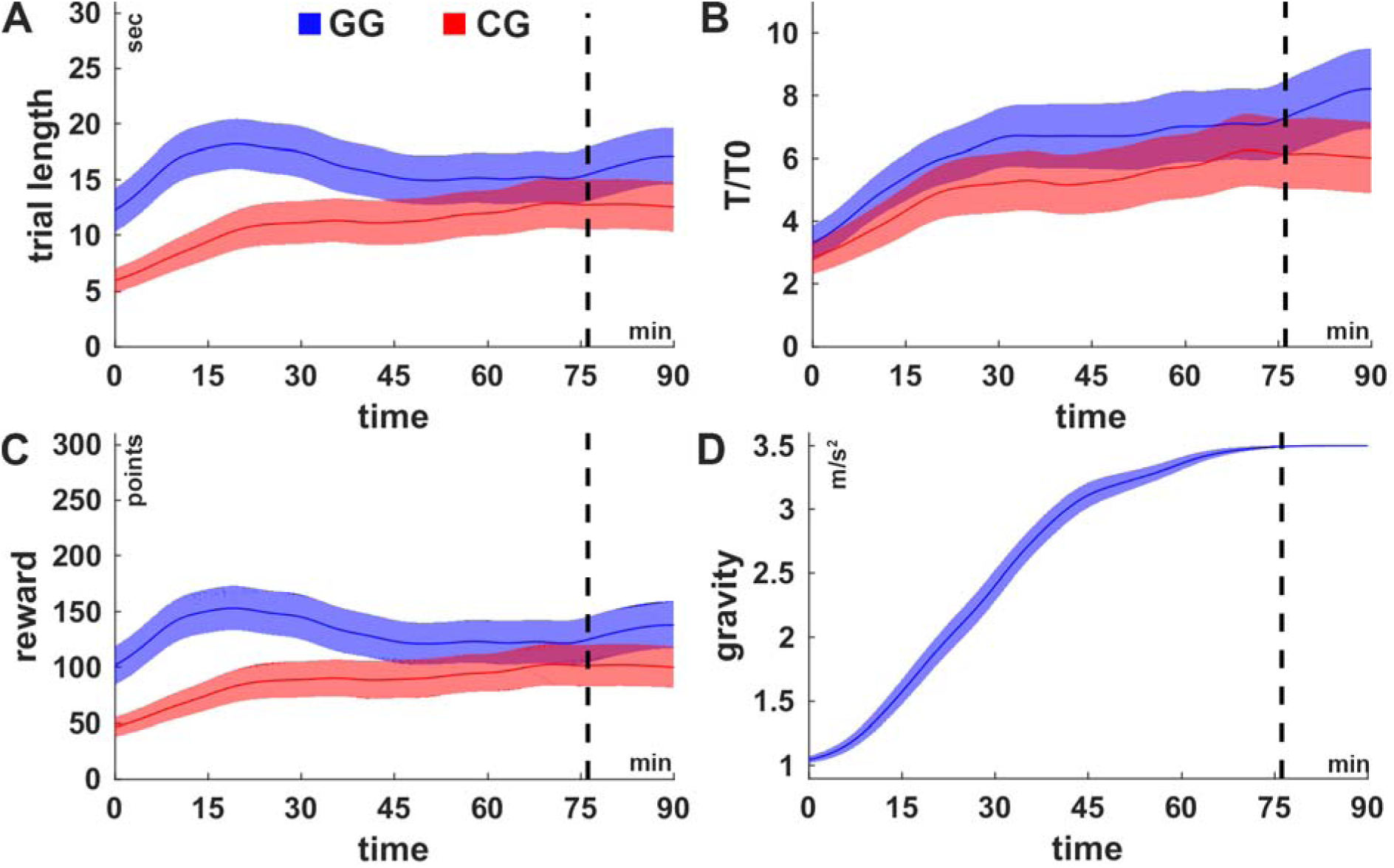
Learning curves. (A) Trial length, (B) normalized trial length (T/T_0_), (C) cumulated reward per trial across the experimental duration in the conditions GG (blue) and CG (red). The normalized trial length (B) reveals a monotonic improvement for both groups whereas the other measures (trial length and reward) are non-monotonic. (D) The average increase of the gravity of subject in condition GG. The shaded areas indicate the inter-subject-variability (±1SEM). The black dashed lines indicate the time at which all subjects in condition GG have reached maximum gravity (g_max_=3.5m/s^2^) latest. For the purpose of illustration, the curves were smoothed over time.

We found no significant difference between the groups in T/T_0_ over the first 5 minutes (Wilcoxon rank sum, **p=0.15**), which suggests that the difference found for the trial length is mostly because of the difference in difficulty and is successfully cancelled out by the normalization. The normalization had no effect on the group difference at the end of the experiment, meaning that subjects in the gradual group are, also according to this measure, significantly better (Wilcoxon rank sum, **p<0.05**). Thus, neither age, gender nor gravity explain the group difference at the end of the experiment.

### Success & failure

We therefore analysed another important incentive for improvement in this task, which is reward. The cumulative reward at the end of the trial is shown in Fig 2C. However, we found that the cumulative reward is mainly determined by the trial length, and does therefore not provide further insight.

In comparison to numeric reward, success (and failure) in balancing is a very strong reward signal. Subjects, who manage to balance the system early during the experiment, have more time to practice and reinforce the successful behaviour. Subjects in condition GG were significantly earlier able to balance the system for the first time (time of first successful trial, Wilcoxon rank sum, **p<0.001, median: GG=2.8min, CG=24.2min**) and were thereby able to practice and explore successful behaviour more deliberately in comparison to subjects in condition CG.

The time between two successful trials (inter-success intervals) is yet another important factor for reinforcement of behaviour. Regression analysis using a linear mixed-effects model was conducted to examine the effect of group and index of success on the (log) inter-success interval (Fig 3). Both factors as well as their interaction were significant (all **p<0.001**). Post-hoc comparison of the groups however did not reveal any significant difference (**p=0.16**), when the model was fitted over the first 35 success. But after constraining the model to the first 20 successes, both factors and their interaction reached significance (all **p<0.001**) and, additionally, post-hoc comparison of the groups revealed a significant difference (**p<0.01**). This result shows that, at the beginning of the experiment, subjects in condition GG receive positive reward (in form of success) more frequently than in condition CG.

Correlation analysis within the groups has revealed a significant relation (Spearman’s correlation coefficient, **GG: rho=-0.57, p<0.05; CG: rho=-0.59, p<0.01)** between the average inter-success internal over the first ten intervals and the task performance at the end of the experiment (average trial length over last 5 minutes). Performing the correlation analysis across both groups revealed the same relation (Spearman’s correlation coefficient, **rho=-0.71, p<0.001**). Overall, these results emphasize the importance of early and frequent success for reaching high performance towards the end.

**Fig 3.**
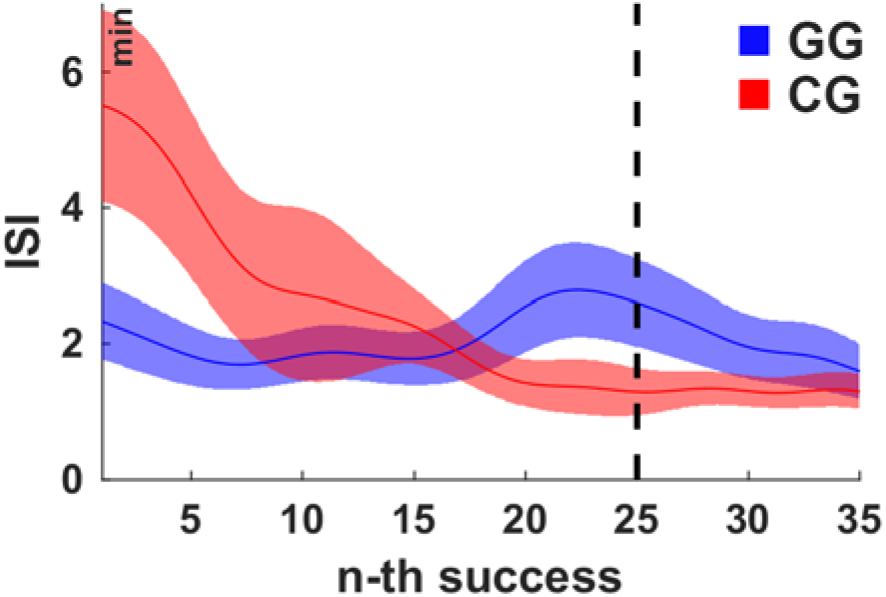
The average inter-success intervals for the first 35 successes. The inter-success intervals (ISI) in condition CG decrease monotonically whereas in condition GG an intermediate increase is observable just before subjects reached the maximum gravity after 25 successes. The shaded areas indicate the inter-subject-variability (±1 SEM). The black dashed line indicates the time at which all subjects in condition GG have reached maximum gravity (g_max_=3.5m/s^2^) latest. For the purpose of illustration, the curves were smoothed over time.

### Timing and Variability of Actions

Analysis of the action timing over the course of the experiment (see Fig 4A, and section Methods for detailed description of temporal measures) revealed a monotonic change to earlier (more predictive) performance of actions (Spearman’s correlation coefficient, **GG: rho=-0.55, p<0.001; CG: rho=-0.15,p<0.05**). Similarly, correlation analysis revealed a significant decrease in action variability (Fig 4B) over the course of the experiment in both conditions (Spearman’s correlation coefficient, **GG: rho=-0.19, p<0.002; CG: rho=-0.19, p<0.002**). This result shows that improvement in action timing is part of the skill acquisition, which will be discussed further in the context of predictive control and the use of internal forward models (see Discussion).

### Gravity-dependent analysis of the performance and actions

Analysis of the average trial length (Fig 5A) across and within gravity steps revealed significant improvement during phases of constant gravity (〈P_2,g_ - *P*_1,g_〉_g_, t-test, **p<0.001**) and significant deterioration after an increment in gravity within the gravity steps (〈*P*_1,g+1_ - *P*_2,g_〉_g_,t-test,**p<0.001**). Furthermore, we found that the action timing changes (Fig 5B) towards an earlier (more predictive) execution of counteractive actions during phases of constant gravity (〈AT_2,g_ - AT_1,g_〉_g_, t-test, **p<0.001**). Coherently with the changes in performance, the action timing is deteriorated after a change in gravity, i.e. actions are timed later, meaning rather in reaction to the events than predictive (〈AT_1,g+1_ - AT_2,g_〉_g_, t-test, **p<0.001**). In alignment with these results, the action variability (Fig 5C) decreases significantly during phases of constant gravity (t-test, **p<0.05**) and increases after an increment in the gravity (for g**>**1.3, t-test, **p<0.05**). These results indicate that the control policy is continuously adapted to the repetitively changing system dynamics during the condition GG.

**Fig 4.**
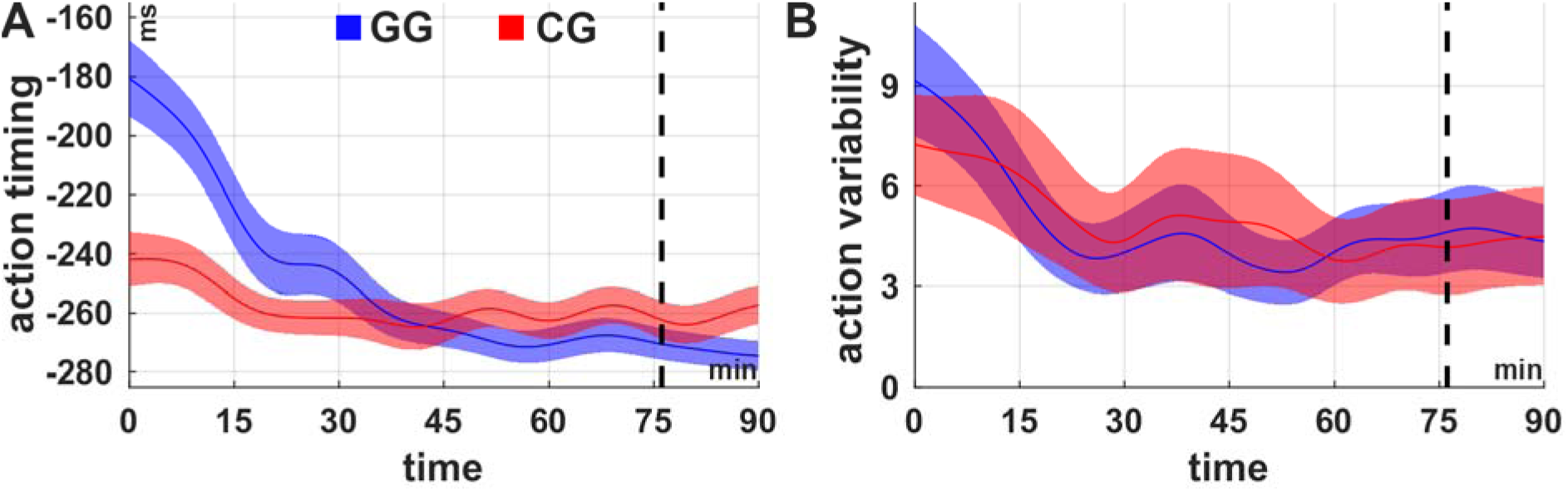
Action timing and variability over the course of learning. Average (A) action timing and action variability over the course of the experiment for the conditions GG (blue) and CG (red). In both conditions the action timing as well as the action variability decline (actions are timed earlier, variability decreases) over time. Both measures were calculated in bins of 5 minutes length. The shaded areas indicate the inter-subject-variability (±1 SEM). The shaded areas indicate the inter-subject-variability (±1 SEM). The black dashed lines indicate the time at which all subjects in condition GG have reached maximum gravity (g_max_=3.5m/s^2^) latest. For the purpose of illustration, the curves were smoothed over time.

**Fig 5.**
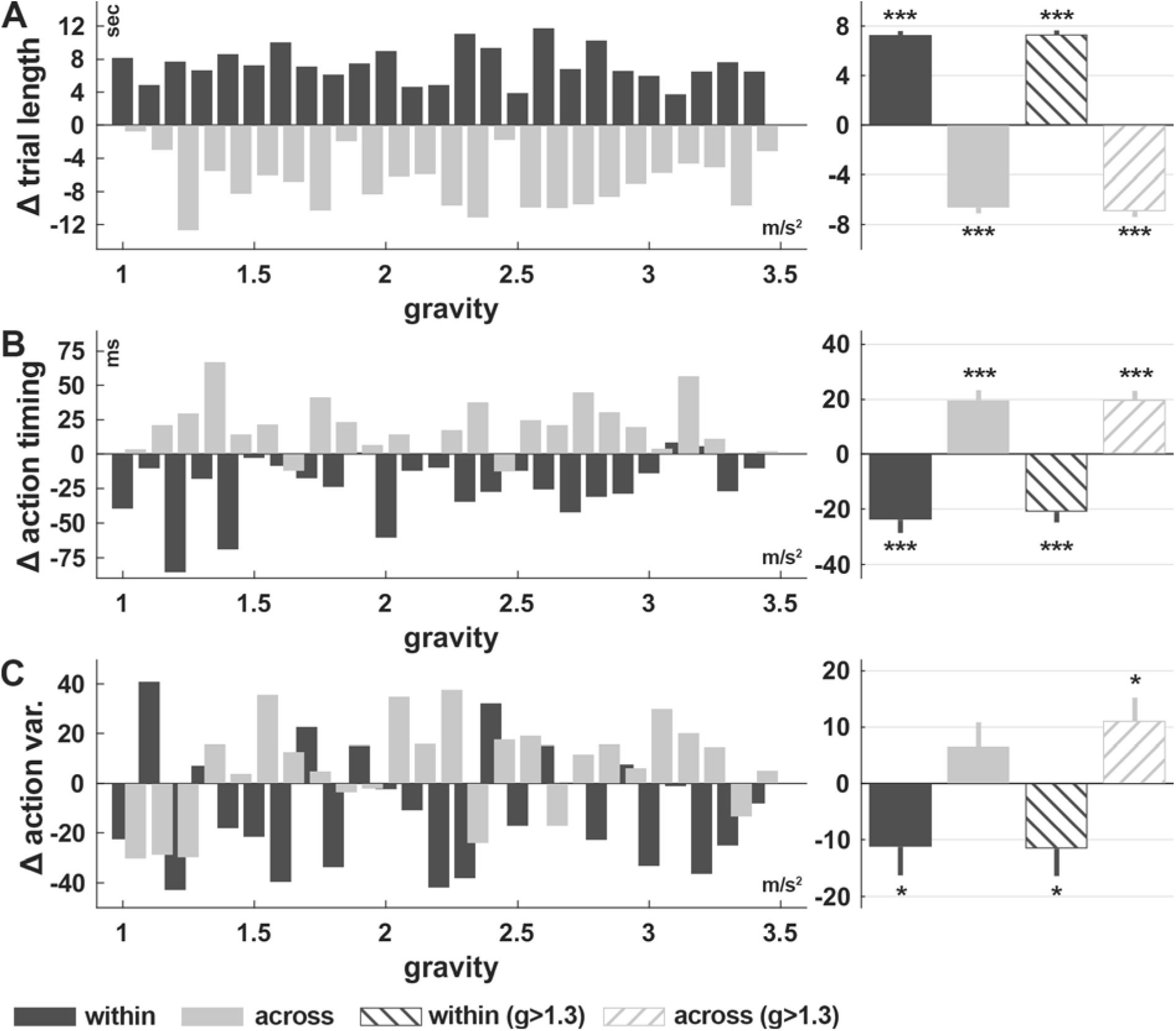
Change of the trial length, action timing and variability as function of the gravity change. (A) Trial length, (B) action timing and (C) action variability. On the left, the measures are shown for each gravity step, on the right as average over all steps is shown with error bars (±1 SEM). Subjects improve (increase in trial length, more predictive actions, less variable actions) within and get worse (decrease in trial length, less predictive actions) across the gravity steps. Excluding the first three gravity steps reveals also for the action variability a significant negative influence of the increments in the gravity. Significance codes: *** p**<**0.001, * p**<**0.05.

### Relationship between Action Timing, Action Variability and Task-Performance

The coherent influence of a change in the gravity on the task performance, action timing and variability suggests a strong relationship, which is independent of learning. After subtracting the coherent influence of learning on the two measures (see Methods and S2 Figure), correlation analysis has revealed a significant relation between the trial length and action timing (Fig 6 A) for both groups (Spearman’s correlation coefficient, **GG: rho=-0.69, p<0.001; CG: rho=-0.50,p<0.001**). Although, we also found a significant correlation between trial length and action variability (Fig 6B) for the condition CG, the relation seems to be much weaker (Spearman’s correlation coefficient, **GG: rho=0.07, p=0.50; CG: rho=-0.22, p<0.05**). In summary, high task performance relies on appropriate action timing.

**Fig 6.**
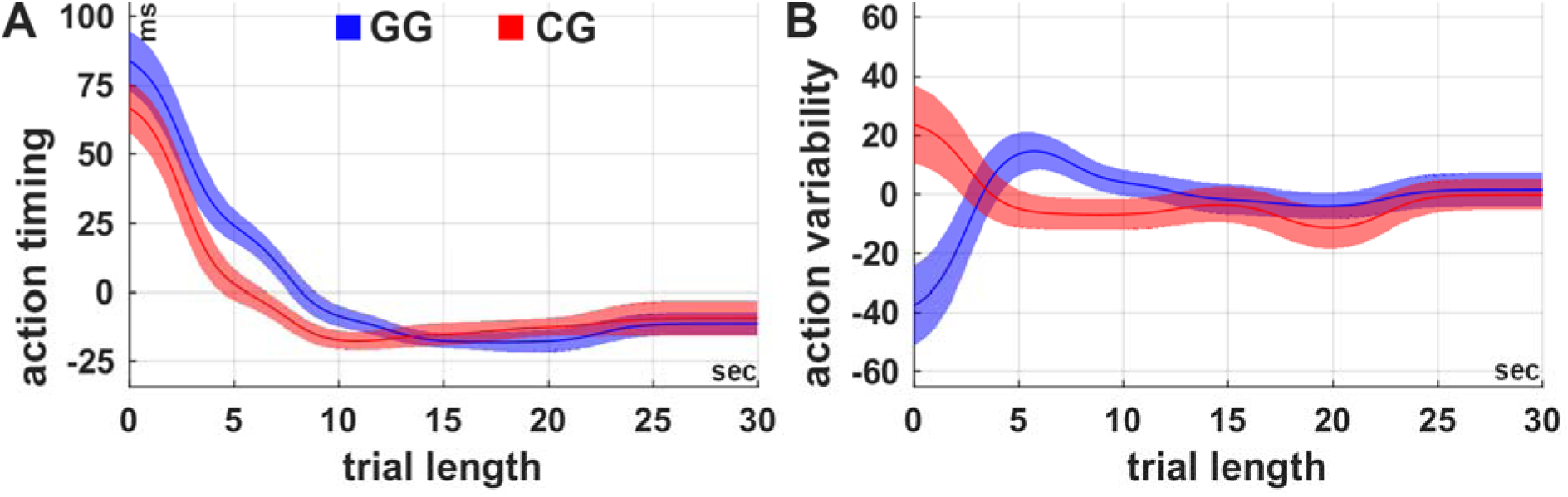
Relationship between task performance, action timing and action variability. Action timing and action variability were normalized with respect to learning (see Methods) and can therefore take negative values. (A) In both groups, there is a strong relationship between the normalized action timing and trial length (task performance). Generally, the more predictive actions are performed (negative action timing) the better the task performance. (B) Except for the low task performances (trial length shorter than 5 seconds), the action variability is not related to task performance. The shaded areas indicate the inter-subject variability (±1 SEM). For the purpose of illustration, the curves were smoothed over time.

## DISCUSSION

The aim of this study was to elucidate how reward-based and error-based learning interact in one single task. We therefore investigated subjects’ learning behaviour in the cart-pole balancing task, which combines skill acquisition and internal model adaptation due to gradually changing dynamics of the cart-pole system. As hypothesized, we have found three main effects: (1) gradually increasing the difficulty of a task, increases the amount of reward subjects receive early in the experiment and is therefore beneficial for acquiring a skill. (2) When these changes influence the controlled system dynamics, there is not only a temporary decrease in general performance but also a specific degradation in the timing and variability of actions. Finally (3), performing actions predictively is crucial for successful the cart-pole balancing.

### Gradual task-difficulty facilitates skill acquisition by means of success

Consistent with many studies on skill acquisition from a wide range of disciplines, we found that gradually increasing the difficulty of a task is beneficial for acquiring a motor skill. Choosing an adequate initial task complexity and increasing the complexity in relation to the improvement during learning, is seen as important prerequisite for efficient motor learning and is, thus, commonly used as training principle in sports as well as in rehabilitation^16,45.^

The theoretical relationship between task difficulty, skill level and learning was systematically described in the “challenge point” framework^46^. The optimal challenge point characterizes the optimal task difficulty, for learning, as function of skill. Guadagnoli and Lee^46^ distinguished between the (a) nominal and (b) functional task difficulty. While the nominal task difficulty is a feature of the task, the functional task difficulty depends in addition to the task also on the skill level of the learner. As part of their framework, they relate the task difficulty and skill to the amount and ability to process information. They elaborate that learning is inefficient if the amount of information is too high (i.e. functional task difficulty is too high), or the capability to process the available information is too low (i.e. skill level is too low). Thus, regarding the efficiency of learning, there is an optimal rate of increasing the task difficulty, which depends on the learner’s skill level.

Consistent with this framework, novices benefited, in our experiment, from an initially low gravity level, yielding low time-constraints for the control of the pole (see S2 Figure). By increasing the gravity after every success, we increased the task difficulty in relation to the skill level and maintained the challenge of the task. In particular, by tightening the time-constraints we gradually encouraged the use of predictive control, which was also identified as crucial mechanism for successfully controlling the cart-pole system.

Despite these more theoretical frameworks and observational studies, mainly in sport science, there is a limited number of studies investigating the effects of adaptively increasing the task complexity on motor learning processes. Most related, is a study by Choi et al.^38^ in which the authors examined retention of different visuomotor transformations over 4 days. They found that the retention is largely improved by adapting the number of trials as a function of performance in comparison to maintaining fixed random scheduling. Varying the difficulty, in terms of limiting the maximum movement time, as a function of performance also improved learning, although to a lesser extent. Since further studies on this topic are lacking, an interesting and still open research question is how the task difficulty has to be increased in order to maintain continuous optimal challenge. Especially for tasks like ours, in which the increase of task difficulty also results in a change of object dynamics, this is an intriguing question. In this case, optimal training schedules have also to take into account that adaptation to changes in object dynamics are necessary. Fewer increments in gravity imply fewer adaptations but at the cost of large increments in task complexity, which might influence reward and motivation negatively.

### Internal forward models support the predictive control of object dynamics

In addition to the substantial amount of studies examining internal model adaptation in classical visuomotor adaptation paradigms, several studies have also shown that humans learn and maintain internal models of object dynamics for optimizing their actions and the corresponding sensory consequences^3,29,30,47–50^. For balancing an inverted pendulum, for example, Mehta and Schaal^30^ have demonstrated that actions of trained subjects performed during short absence of visual feedback (450-550ms) do not significantly differ from actions with visual feedback. They suggested that predictions formed by an internal forward model of the cart-pole system replace the visual feedback during these phases. Our results are in line with this hypothesis and imply predictive control as a critical mechanism for successfully mastering the cart pole task. In our experiment, subjects performed the actions progressively in advance in relation to the state of the system (Fig 4A), which indicates gradually improved predictive behaviour. Emphasizing the importance of action timing even further, we demonstrated a crucial relationship between the measured action timing and cart-pole balancing performance (Fig 6A).

### The adaptation of forward models during the gradual increase of gravity

In order to profit most from internal forward models of object dynamics, they have to be acquired in the beginning of the learning process and need to be adapted when the behaviour of the modelled object changes^32^. Intriguingly, when interacting with novel objects, people seem to learn to predict the behaviour of the objects before they can master its control47. For real balancing of a pole on the fingertip, Lee and colleagues^44^ suggested that subjects acquire an accurate pendulum model, which accounts for the gravitational dynamics and mass distribution along the pole, already within the first moments of wielding the pole through “dynamic touch”^51^.

On the other hand, changes to the object dynamics, such as increments in gravity in our experiment, result in inaccurate extrapolations of the system’s state, which primarily leads to deteriorated performance (Fig 5A). A more specific prediction is that the acceleration of the pole is underestimated after an increase of the gravity. Thus, with respect to a certain pole angle, responses would be delayed. Our analysis of the change in action timing across the gravity steps (〈AT_1,g+1_ - AT_2,g_〉_g_) supports this hypothesis (Fig 5B). Consequently, the altered system dynamics and the now imprecise predictions of the forward model lead to increased spatio-temporal variability (Fig 5C). These observations are consistent with previous studies showing temporarily increased spatio-temporal variability in virtual bouncing game after changing gravity^52^. As soon as the internal model is adapted to the new dynamics of the object, state estimates are again accurate, leading to normal motor variability and performance.

### Gradual increase of difficulty in skill acquisition tasks and sensory-motor adaptation paradigms

In recent years, increasing attention has been gained by sensorimotor adaptation studies which examine the effects of perturbing the sensorimotor mapping of reaching gradually instead of suddenly^8,9,53,54^. The (performance-independent) gradual perturbation schedule does not predominantly facilitate learning, but leads to an increased and prolonged after-effect after removing the perturbation ^8^. It has been suggested that gradual perturbation schedules favour implicit learning mechanisms (involving internal forward models) in contrast to explicit rule-based learning^53^.

Even more influential are gradual perturbation paradigms for sensorimotor adaptation when binary reward-based feedback is provided instead of end-point errors in reaching. In such paradigms only hitting the target is positively rewarded, which is difficult if perturbations are large and sudden^10–12^. Thus, finding the right correction for the perturbation and receiving positive reward is difficult, which makes reward-based learning tough. In contrast, if perturbations are gradual, the necessary adaptation between subsequent perturbation steps requires less exploration and may even be in the order of magnitude as subjects’ motor variability^55,56^, leading to implicit learning and fast adaptation of the underlying internal forward models. From a computational point of view, instead of searching for an adequate solution, which is in the sensory-action space faraway, the gradual perturbation paradigm allows for guided exploration in small steps with intermittent reward.

This view is transferable to our task when task complexity is gradually increased. Based on simple heuristic control rules, which are sufficient to solve the problem (balance the pole) under the easy initial task condition of low gravity (g_0_=1.0m/s^2^), the exploration during learning is guided stepwise through the sensory-action space to a solution for the complex task condition (g_max_=3.5 m/s^2^). The potential benefits of this approach are supported from previous results from machine learning, showing that the use of a fuzzy controller to implement heuristic knowledge and reward-based learning mechanisms for the refinement of control strategies can facilitate the learning processes substantially^57^. By guiding the exploration of the sensory-action space, subjects receive primarily more frequently positive reward, especially early in the learning process.

Success and reward have shown to be strong reinforcement signals, which can not only accelerate learning^12,58^ but also have a motivational influence on the subject^59,60^. Moreover, it has been shown that offline gains and long-term retention of newly formed motor memories benefit from training under rewarding conditions^36^, demonstrating the power of reward. Recently, it was shown by Therrien et al.^11^, that adaptation to sudden sensorimotor perturbations is possible with binary reward, if the reward is provided relative to the current mean performance (closed loop reinforcement). Hence, instead of providing positive reward only if the target was hit, reward is used to guide the exploration towards the target, by providing positive reward if the action was better than the previous ones. Thus, instead of changing the task demands, the reward landscape is shaped by the current performance. Both approaches demonstrate the benefit of incorporating the functional task difficulty^46^, which is set by the current skill level of the learner, into the design of training schedules.

## CONCLUSIONS AND OUTLOOK

In this study, we have presented an experimental setup to investigate reward-based motor skill acquisition to control an external object with changing dynamics.

We conclude that gradual increase in task difficulty accompanied with changes in object dynamics facilitates, despite interleaved brief degradations in performance, the skill acquisition by means of success-mediated learning. Interleaved degradation in performances due to changes in object dynamics is associated with less predictive timing and increased variability of actions. The presented results motivate several further studies, in order to examine the interplay between reward-based skill acquisition and adaptation of the internal model to changes of the object dynamics in more detail. As interesting open questions remain for instance (i) potential after-effects in timing after the gravity is changed back to a lower value, (ii) differences in retention between gradual and sudden increase of gravity as well as (iii) finding an optimal schedule to increase the gravity, in order to minimize the number of changes while preserving the advantage of slowly increasing the complexity. Future studies will have to examine the neural mechanisms of skill acquisition when task difficulty and object dynamics are gradually changed. In particular, the cerebellum, with its role in reward-based learning^61^ and maintenance of internal models of object dynamics, might be an interesting target of further investigation. In conclusion, these studies will advance our knowledge in skill acquisition for complex movements, applicable in several disciplines, such as sports, professional skill development, and neuro rehabilitation.

## METHODS

### Subjects

We analysed thirty right-handed subjects (age range 18-30 years, mean age 23.4; 15 females, 15 males).

All subjects gave informed written consent prior to participation. The study had been approved by the local institutional ethical review board in Tübingen (AZ 409/2014BO2). Subjects were randomly assigned to one of two groups corresponding to the two examined experimental conditions (gradual gravity, GG; constant gravity, CG; see Learning Paradigm). Both groups consisted of 15 subjects with similar average age (GG: 24.1 years, CG: 22.6 years). Gender was balanced between groups (GG: 9 males, 6 females, CG: 7 males, 8 females). All subjects were right-handed and used their right hand throughout the experiment. Subjects’ were reimbursed independent of performance.

### Details of Experimental Setup

The cart-pole system consists of a cart to which a one-meter long pole is attached. Due to the assigned masses (pole: 0.08kg, cart: 0.4kg) gravity forces the pole to rotate downwards when not being perfectly upright. We did not simulate friction. Forces from the left or right are applied by the subjects to the cart in order to control the system. Participants controlled the virtual force using a SpaceMouse® Pro (3Dconnexion, Fig 1B). This input device is comparable to a joystick but it can measure six degrees of freedom (DOF) including the left-right translation. We used this DOF to control the force. To this end, the lateral displacement of the device’s knob (±1.5mm) relative to the rest position in the centre of the device was translated into a virtual force into the same direction with proportional magnitude that pushes against the cart. Hence, a rightward knob movement causes a virtual force, which pushes the cart to the right. The knob of the device is automatically pulled back to the rest position in the centre of the device (the device exerts a force of 7.4N at full lateral displacement). The simulation of the cart-pole dynamics was implemented in MATLAB (The MathWorks, Inc.) using the 4^th^-order Runge-Kutta method. Visual feedback was provided on a 15 inch monitor using the Psychtoolbox^62^–^64^ at a refresh rate of 60Hz. Correspondingly, the time-discretization constant Δt of the simulation was set to 1/60s. Subjects were not constrained in posture but were asked to sit comfortable about 60cm away from the monitor. The input-device was aligned with the monitor such that the left-right knob movement was in correspondence with the virtual force and cart movement on the monitor.

### Learning Paradigm and Experimental Conditions

At the beginning of every trial the cart-pole system is initialized with a random pole angle drawn uniformly from [-7.5, 7.5] degrees, positioned at the center of the track with both velocities (cart and angular pole velocity) set to zero.

The pole has to remain within the green circular segment (±60 degree, Fig 1A) while the cart must not leave the track (±5m). This has to be achieved by applying forces of up to 4N from either side to the cart using the input device. The forces accelerate the cart, depending on the direction, to the left or right by which the pole can be balanced. A trial was considered successful, if balance was maintained for 30 seconds without violating any constraint. Hence, trials were at maximum30 seconds long. Violation of one of the constraints (cart position, pole angle) also terminates the trial. The number of trials was not limited, instead we limited the duration of the experiment (see below). Before the next trial begins, feedback about the violated constraint and the duration of the trial is provided.

In addition to the terminal feedback at the end of every trial, subjects also receive cumulative reward during the trial, which is displayed as number in the cart (Fig 1A). The theoretically maximum reward per second is 10 points, which can be achieved by holding the pole perfectly vertical, keeping the cart exactly in the centre of the track while not applying any force to the cart (for details, see S1 Appendix). Thus, in theory, a maximum reward of 300 points per trial is achievable. This ultimately means that subject has to keep the system within the constraints for 30 seconds, i.e. balance the pole on the cart for 30 seconds without leaving the track. Subjects were instructed about these factors and were asked to maximize the reward in every single trial.

### Data Processing and Analysis

We analysed the trial length, reward per second and cumulative reward of every trial (including the unsuccessful balancing attempts) in order to quantify improvement. The trial length T can be characterized by two factors: (i) T_0_, which is the time it takes for the pole to violate the pole angle constraint when no controlling input would be present (S1 Figure) and, (ii) the time by which the trial is prolonged due to subject’s actions. The time T_0_ is completely described by the initial pole angle and the gravity. In a low-gravity environment (e.g. at the beginning of condition GG) the time T_0_ is much longer than for higher gravity values (e.g. throughout condition CG or at the end in condition GG). Hence, in condition GG T_0_ decreases as function of the increasing gravity during the experiment and progressively limits the time in which subjects have to counteract (tightening of time-constraint). We simulated the system with different initial pole angles and thereby revealed a nonlinear relation between T_0_ and the gravity (S1 Figure). In order to account for this factor, we determined T_0_ for every recorded trial and calculated the ratio T/T_0_, which is a measure of how much better (**>**1) or worse (**<**1) the subject was controlling the system compared to doing nothing.

Other important factors, especially in the context of reward-based learning, are reward, success and the time passed between subsequent successes. In the following, we call the latter inter-success intervals (ISI).

### Determining the action timing and variability

In order to infer more about the underlying mechanisms of improvement, we quantified the changes in subjects’ control policies. We therefore examined the applied forces as function of the system state, specifically as function of the pole angle (Fig 7A). The rationale of our approach is to define events in the state space and analyse the applied forces relative to those events (event-triggered averaging, Fig 7B & C). We focus our analysis on the situations when the pole is tilted by a certain angle and is rotating downwards. In these situations, which we describe as events, a counter-action is necessary. Using our method we determined when and how variable these counter-actions were performed. The following four steps describe the procedure in more detail.

**Fig 7.**
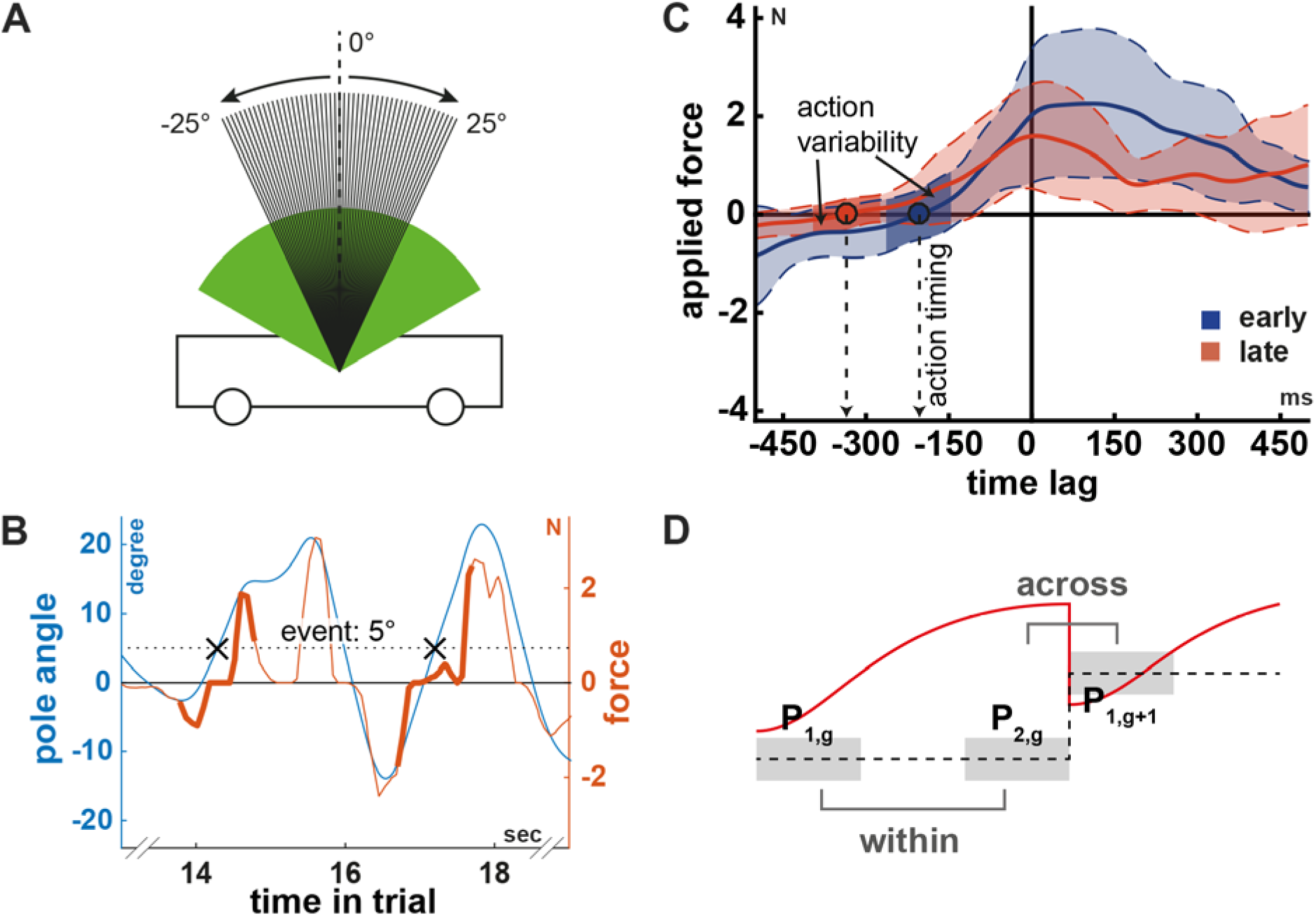
Schematic of the action timing and the change of measures as function of the gravity. (A) All pole angles investigated as events (integer valued pole angles from -25 to 25°). The arrows indicate the direction of the pole movement. (B) Pole angle (blue) and input force (orange) trajectories in a representative trial illustrating two event occurrences (black crosses) and corresponding two force segments (thick lines). Negative force values correspond to a leftwards force. Aligning all segments that correspond to one event and averaging the segments of different trials in a window of 2 minutes yields the curves shown in panel C. (C) Average force segments during two periods (early: blue, late: orange) in learning of a representative subject for illustration of the action timing and variability measures. The circles indicate the time (action timing) when the subjects changed the direction of the force relative to the occurrence of the event (zero time lag). Positive time lags describe the time after the event occurred. As expected, actions early (blue) during learning are made rather in reaction to event occurrences (towards positive time lags) whereas learning leads to the ability to make actions predictively (more negative time lag). Coloured areas illustrates the variability in force segments. The dark coloured areas indicate the action variability. It is the average standard deviation of the force segments ±60ms around the zero crossing (action timing). The variance in input force round the zero crossing (action variability) is lower for actions late during learning, suggesting more consistency. (E)Illustration of the procedure to examine changes in different measures relative to an increase in gravity. We here show exemplarily the trial length (red curve) in relation to the gravity (dashed line). The light grey areas illustrate the periods under investigation. Subtracting the average trial length in the highlighted periods yields the change within (*P*_2,g_-*P*_1,g_) and across (*P*_1,g+1_-*P*_2,g_) the gravity step(s).

i. We defined the integer valued pole angles in the range from -25 to 25 degrees as events (Fig 7 A). For each of these 51 events we determined the occurrences in every trial. Event occurrences between two time-discretization steps (frames) were estimated using linear interpolation.
ii. We then excluded all event occurrences in which the pole is actually rotating upwards and therefore no counter-action is required. We further excluded extreme pole angle velocities. To this end, we excluded all event occurrences in a running window of two minutes across trials, for which the pole angle velocity did not lay between the 20%-and 80%-quantiles of all observed pole angle velocities.
iii. Next, we extracted segments of one-second length from the force input trajectory, which are centred on the previously determined event occurrences (Fig 7B). These segments look roughly like sigmoidal functions going from negative to positive force values or vice versa, corresponding to a left-right or right-left movement of the device knob by the subject.
iv. In the last step, we averaged all segments corresponding to one event (pole angle) in a running window of 2 minutes length across trials and determined the zero crossing of the average segment (Fig 7C). Thereby we find the time of change from a negative (leftwards) to a positive (rightwards) force (or vice versa) relative to the event occurrence (zero time lag).

We refer to this time (when the actions changes relative to the state of the system) as action timing. Notice, that we do not interpret this measure as the reaction time, even though it might be related. Furthermore, we estimated the variability of the actions by calculating the mean standard deviation of the applied forces in a centred window of 120ms length around the zero crossing (Fig 7C).

### Relationship between action timing and task performance

During the analysis of the influence of the action timing and variability on the task performance, we faced the problem that all three measures change over time due to learning. Correlation analysis between these measures therefore trivially revealed high correlation. We were however interested in verifying the action timing and variability as crucial measures for achieving high balancing performance. Hence, we were looking for a normalization, which eliminates the concurrent influence of learning on all measures but preserves potentially existing correlations between the measures. Since we did not measure learning per se, we had to use time as representative. In order to express the action timing and action variability as functions of task performance (trial length) and time, we discretized trial length and time (S2 Figure A). Trial length was split into bins of 5 seconds, while time was split into bins of 5 minutes. Within each of the two-dimensional bins, the action timing and action variability have been determined. We eliminated the influence of learning by subtracting the average action timing (action variability respectively) across all trial length bins within each time bin (S2 Figure A & B). The action timing and variability are thereby expressed as function of the discretized trial length, independent of the influence of learning.

### Quantification of changes induced by gradual increments in gravity

We next analysed the influence of changing the gravitational constant on the performance, action timing and motor variability in the gradual group. The gravitational constant is described by a monotonically increasing stairs function over time (Fig 1C). Consequently, there are phases of constant value and sudden but small changes from one trial to the next (Fig 7D). Let *P*_1,g_ denote the average trial length during the first third of the gravity step with value g and let *P*_2,g_ denote the average trial length during the last third of this step. We estimated the improvement within gravity steps by taking the difference *P*_2,g_ - *P*_1,g_. Averaging these across all steps yields a single value 〈*P*_2,g_ - *P*_1,g_〉_g_ for each subject, representing the average improvement during periods in which the system does not change. Similarly, we estimated the influence of an increment in gravity by calculating the average trial length over the last third (*P*_2,g_) and the first third of the next step (*P*_1,g+1_). The average difference over all gravity steps 〈*P*_1,g+1_ - *P*_2,g_〉_g_ measures the average change in performance caused by the changes of the system dynamics. Notice, that we denote the average of a measure across all gravity steps g by 〈·〉_g_.

For the action timing and variability, we proceeded similarly but calculated the differences for each event (pole angle) separately and then averaged over all events and gravity steps. Thereby we can analyse changes in the action timing (AT) and variability (AV) during periods of constant system dynamics (〈AT_2,g_ - AT_1,g_〉_g_ and〈AV_2,g_ - AV_1,g_〉_g_) and the influence of sudden changes (〈AT_1,g+1_ - AT_2,g_〉_g_ and 〈AV_1,g+1_ - AV_2,g_〉_g_).

### Statistics

All statistical analyses were performed in R (v3.3.2) using the package lme4 (v1.1), lmerTest (v2.0), phia (v0.2). Significance tests for differences between populations were performed using the two-sided t-test if measures were normally distributed according to the Shapiro-Wilk-Test. Otherwise Wilcoxon’s rank-sum test for equal medians has been used. Relationship between measures was evaluated using Pearson’s linear correlation coefficient if measures were normally distributed according to the Shapiro-Wilk-Test. Otherwise Spearman’s correlation coefficient was used. For the analysis of the measures over time, correlation coefficients were computed or linear mixed-effects models were used, which included a random effect for the intercept of each subject.

